## Supplementary Information for "Perception of frequency modulation is mediated by cochlear place coding"

### Supplementary Text

#### *Quantifying contributions from absolute thresholds, age, and sensitivity to AM*

The correlations between the low slope and FM detection demonstrate a significant role for frequency selectivity, even after accounting for sensitivity to AM, age, and absolute thresholds. But this analysis does not address the question of how much each factor contributes to FM sensitivity. Unlike correlations, which are bi-directional, multiple linear regression (MLR) is a conservative, directional approach to examining the amount of variance accounted for by each variable. Because many of the variables are correlated, the order the variables are entered into the model will affect the percentage of variance explained by each variable. We took the most conservative approach by entering the low and high slopes last, after all the other variables. Factors known or believed to contribute to FM sensitivity (absolute thresholds in the measured ear, age, sensitivity to AM at the corresponding rate, low slope, and high slope, entered in this order) were entered into the model, fitted using the Ordinary Least Squares method. The full models, with all variables entered, explained 59.5% ($p < .0001$) and 52.1% ($p < .0001$) of the variance in slow and fast FM, respectively. When sequentially entering each variable, absolute thresholds accounted for 45.2% ($p = .022$) and 21.7% ($p = .11$, n.s.) of the variance for slow and fast FM, respectively (note that all of the $p$ values here correspond to the significance of the variable in the full model). Because age is known to impair FM detection (He et al., 2007; Strelcyk and Dau, 2009; Grose and Mamo, 2012; Paraouty et al., 2016; Wallaert et al., 2016; Paraouty and Lorenzi, 2017; Whiteford et al., 2017), age was entered into the model second, accounting for an additional 4.03% (slow FM: $p = .074$, n.s.) and 3.78% (fast FM: $p = .16$, n.s.) of the variance, while AM, entered third accounted for 4.03% (slow FM: $p = .039$) and 18.7% (fast FM: $p < .0001$). The low slope (slow FM: 6.2%, $p = .009$; fast FM: 7.91%, $p = .008$) but not the high slope (slow FM: .03%, $p = .864$; fast FM: .01%, $p = .938$) significantly contributed to the variance in sensitivity to FM at both rates, consistent with the partial correlation analysis (Fig. 6). Note that entering the slopes first, instead of last, into the regression means that the variance explained is the same as the squared correlations plotted in the left-most two panels of

Fig. 5 (e.g., slow FM and low slope: 39.3%; fast FM and low slope: 22.1%), but the total variance explained in the full models is unaffected. Entering just the low slope and AM at the corresponding rate into the MLRs, which would be consistent with Zwicker's (1956) place model, accounts for 44.9% and 43.7% of the variance for slow and fast FM, respectively, with significant contributions from both the low slope (slow FM: $p < .0001$; fast FM: $p < .0001$) and AM (slow FM: $p = .026$; fast FM: $p < .0001$).

***Results with outlier included (n=56)***

Including the outlier participant with unusually poor FM sensitivity generally did not affect the results (Supplementary Table 1). One exception was that the correlation between slow FM and slow AM detection thresholds was significantly different from the correlation between fast FM and fast AM detection thresholds (Z = 2.13, $p = .032$, two-tailed) when including this participant, but this effect was no longer present with the outlier removed (see "Correlations between FM and AM detection" in main text). Results for the correlations between FM sensitivity and the slopes of the masking patterns before and after controlling for audibility, sensitivity to AM at the corresponding rate, and age are plotted in Supplementary Figs. 2-3.

**Supplementary Table 1.** Results with outlier subject included demonstrate the same trends as the main text.

| Variable X | Variable Y | Statistic |
|---|---|---|
| Absolute thresholds at 1 kHz | Low slopes | $r = -.703$, $p < .0001$ |
| Absolute thresholds at 1 kHz | High slopes | $r = -.707$, $p < .0001$ |
| Slow AM | Fast AM | $t_{55} = 16.7$, $p < .0001$, $d_z = 2.25$ |
| Slow FM | Fast FM | $t_{55} = -2.31$, $p = .012$, $d_z = -.312$ |
| Slow FM | Fast FM | $r = .867$, $p < .0001$ |
| Slow AM | Fast AM | $r = .617$, $p < .0001$ |
| Fast FM | Fast AM | $r = .45$, $p = .0006$ |
| Slow FM | Slow AM | $r = .2$, $p = .068$ |

**Supplementary Table 2.** AM incoherence discrimination results.

| **Effect** | **$DF_{effect}$** | **$DF_{error}$** | ***F*** | ***p*** | **$\eta_p^2$** |
|---|---|---|---|---|---|
| Frequency Separation | 1 | 19 | 72.3 | **< .0001** | .792 |
| Center Frequency | 2 | 38 | 38 | **< .0001** | .667 |
| Rate | 1 | 19 | 10.9 | **.004** | .365 |
| Frequency Separation*Center Frequency | 2 | 38 | 11.6 | **.0001** | .379 |
| Frequency Separation*Rate | 1 | 19 | 1.66 | .213 | .08 |
| Center Frequency*Rate | 1.51 | 28.7 | 16.6 | **< .0001** | .466 |
| Frequency Separation*Center Frequency*Rate | 2 | 38 | .841 | .439 | .042 |

**Supplementary Table 3.** Complex AM detection results.

| **Effect** | **$DF_{effect}$** | **$DF_{error}$** | ***F*** | ***p*** | **$\eta_p^2$** |
|---|---|---|---|---|---|
| Phase | 1 | 19 | 9.93 | **.005** | .343 |
| Frequency Separation | 1 | 19 | .964 | .339 | .048 |
| Center Frequency | 2 | 38 | 3.04 | .059 | .138 |
| Rate | 1 | 19 | 20.9 | **.0002** | .524 |
| Phase*Frequency Separation | 1 | 19 | .183 | .673 | .01 |
| Phase*Center Frequency | 2 | 38 | 25.2 | **< .0001** | .571 |
| Frequency Separation*Center Frequency | 2 | 38 | .029 | .972 | .002 |
| Phase*Frequency Separation*Center Frequency | 2 | 38 | 1.85 | .171 | .089 |
| Phase*Rate | 1 | 19 | 4.94 | **.039** | .206 |
| Frequency Separation*Rate | 1 | 19 | .235 | .633 | .012 |
| Phase*Frequency Separation*Rate | 1 | 19 | 2.68 | .118 | .123 |
| Center Frequency* Rate | 2 | 38 | 1.15 | .326 | .057 |
| Phase*Center Frequency*Rate | 2 | 38 | 7.58 | **.002** | .285 |
| Frequency Separation*Center Frequency* Rate | 2 | 38 | .523 | .597 | .027 |
| Phase*Frequency Separation*Center Frequency*Rate | 1.38 | 26.1 | 1.47 | .244 | .072 |

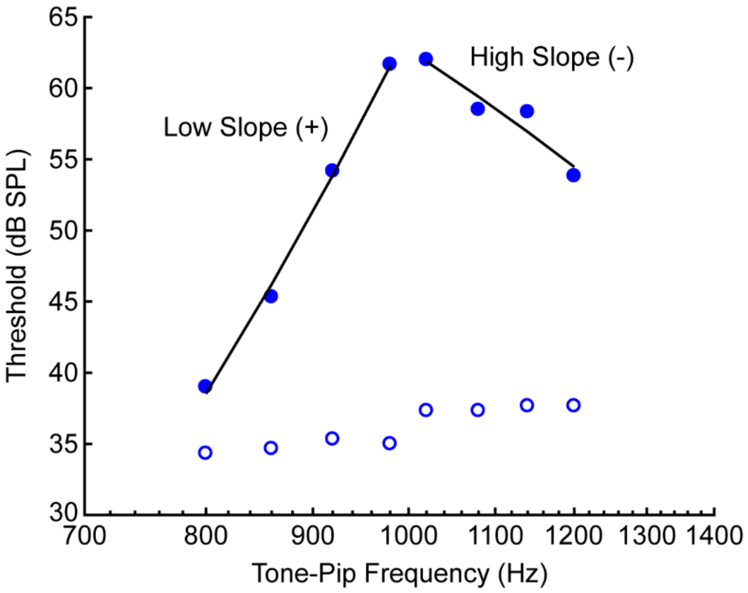


**Supplementary Figure 1.** Example forward masking pattern for a single subject. Absolute thresholds for the 20-ms tone in quiet (unfilled circles) and when preceded by a 500-ms, 1-kHz pure-tone forward masker (filled circles). The level of the tone must be much higher to be perceived when the tone is very close in frequency to the masker as opposed to when it is farther away. The slopes were calculated by conducting two linear regressions: one between the thresholds of the four lowest (low slope) and one between the four highest (high slope) tone-pip frequencies.

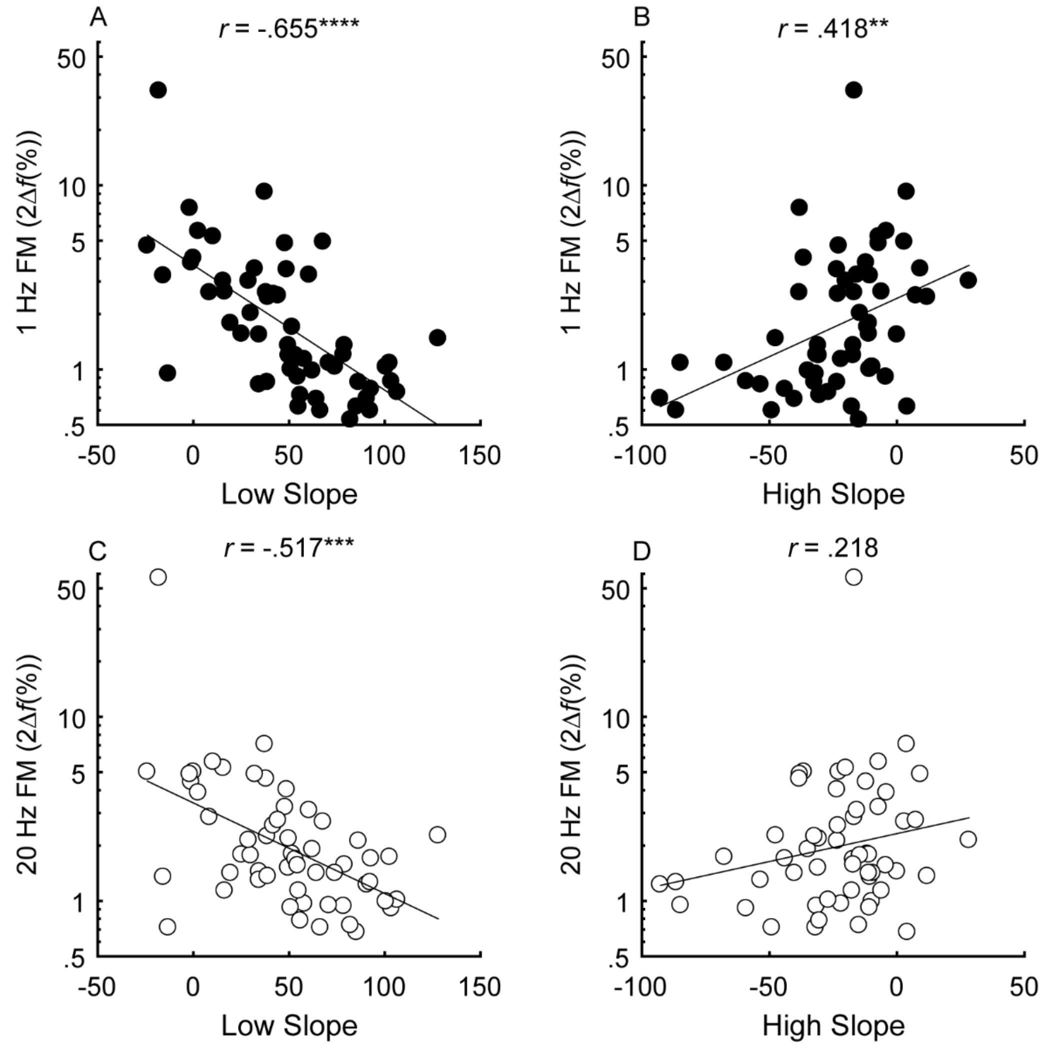


**Supplementary Figure 2.** Correlations between the low slope (A and C) and high slope (B and D) and slow ($f_m$ = 1 Hz; black) and fast ($f_m$ = 20 Hz; white) FM detection (n=56). Correlations marked with an * are significant after Holm's correction (****$p$ < .0001, ***$p$ < .001, **$p$ < .01, and *$p$ < .05).

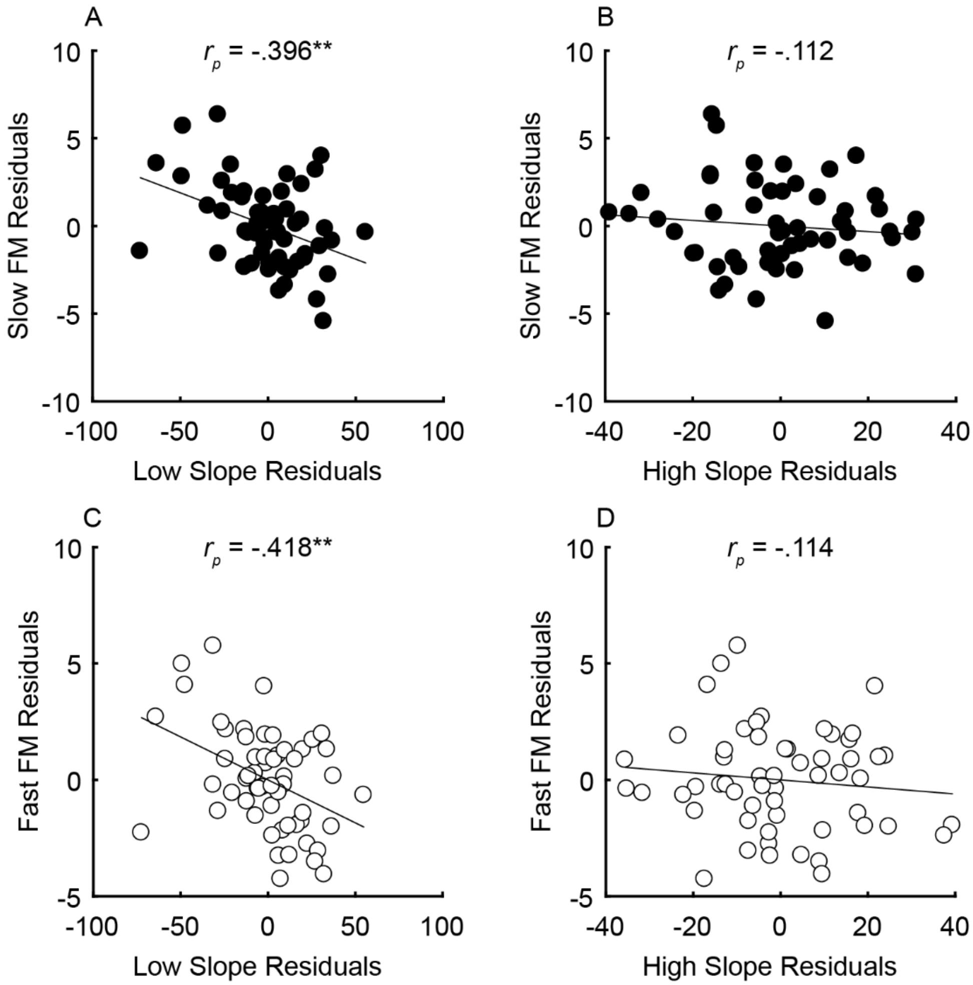


**Supplementary Figure 3.** Partial correlations between the steepness of the masking function slopes (x-axes) and FM detection (y-axes) after variance due to audibility, sensitivity to AM, and age has been partialled out for n=56 participants. Units of the x- and y-axes are arbitrary because they correspond to the residual variance for slow ($f_m$ = 1 Hz; black) and fast FM detection ($f_m$ = 20 Hz; white). Correlations marked with an * are significant after Holm's correction (****$p$ < .0001, ***$p$ < .001, **$p$ < .01, and *$p$ < .05).

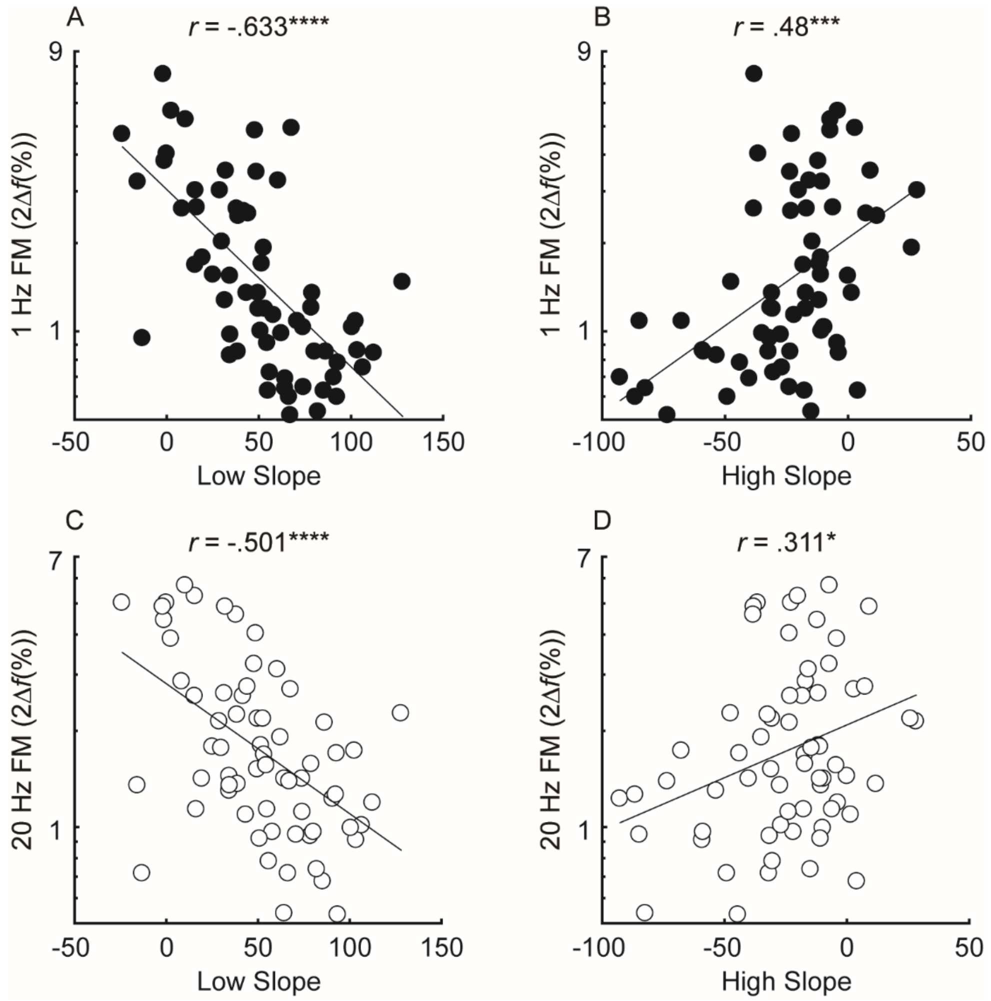


**Supplementary Figure 4.** Correlations between the low slope (A and C) and high slope (B and D) and slow ($f_m$ = 1 Hz; black) and fast ($f_m$ = 20 Hz; white) FM detection, including both ears from the asymmetric subjects (n=66 ears). Correlations marked with an * are significant after Holm's correction (****$p$ < .0001, ***$p$ < .001, **$p$ < .01, and *$p$ < .05).

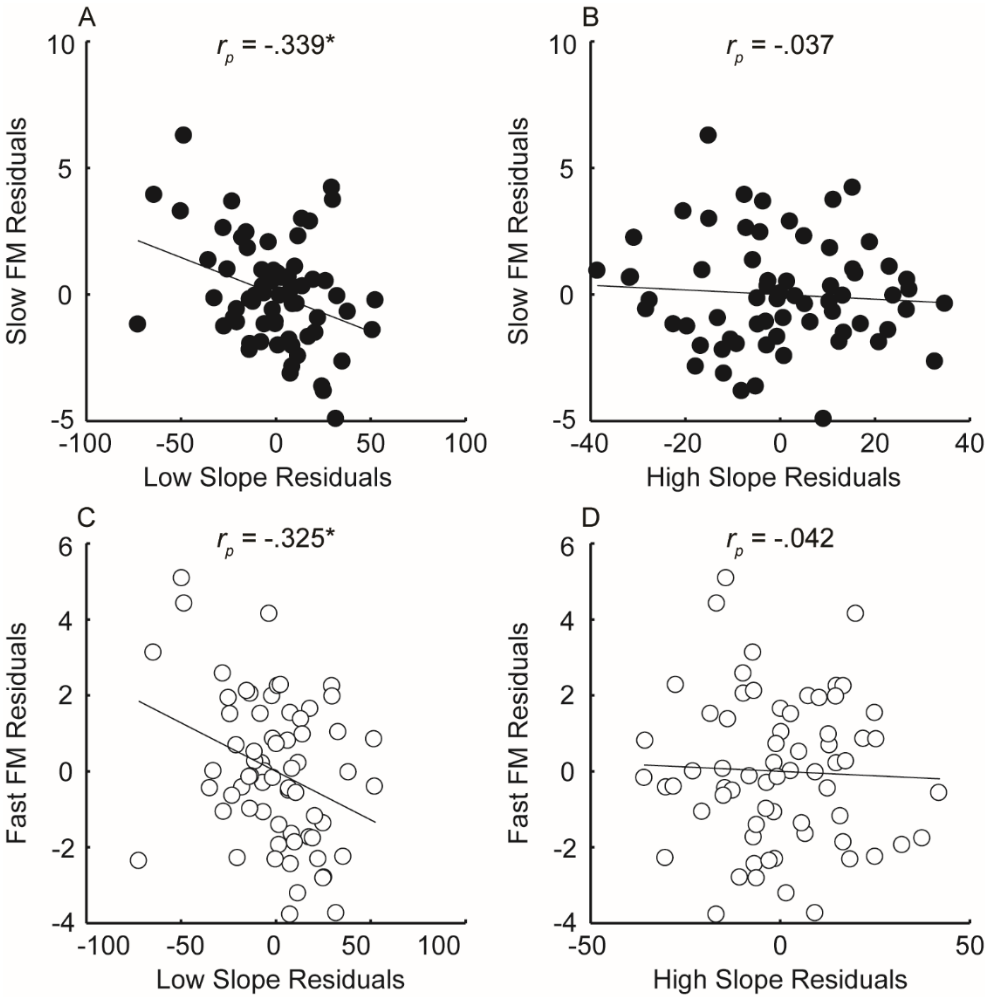


**Supplementary Figure 5.** Partial correlations between the steepness of the masking function slopes (x-axes) and FM detection (y-axes) after variance due to audibility, sensitivity to AM, and age has been partialled out for n=66 ears, including both ears from participants with asymmetric loss. Units of the x- and y-axes are arbitrary because they correspond to the residual variance for slow ($f_m$ = 1 Hz; black) and fast FM detection ($f_m$ = 20 Hz; white). Correlations marked with an * are significant after Holm's correction (****$p$ < .0001, ***$p$ < .001, **$p$ < .01, and *$p$ < .05).
